# A Cut/cohesin axis alters the chromatin landscape to facilitate neuroblast death

**DOI:** 10.1101/299164

**Authors:** Richa Arya, Seda Gyonjyan, Katherine Harding, Tatevik Sarkissian, Ying Li, Lei Zhou, Kristin White

## Abstract

Precise control of cell death in the nervous system is essential for development. Spatial and temporal factors activate the death of Drosophila neural stem cells (neuroblasts) by controlling the transcription of multiple cell death genes through a shared enhancer, enh1. The activity of enh1 is controlled by *abdominalA* and *Notch*, but additional inputs are needed for proper specificity. Here we show that the Cut DNA binding protein is required for neuroblast death, acting downstream of enh1. In the nervous system, Cut promotes an open chromatin conformation in the cell death gene locus, allowing cell death gene expression in response to *abdominalA*. We demonstrate a temporal increase in global H3K27me3 levels in neuroblasts, which is enhanced by *cut* knockdown. Furthermore, *cut* regulates the expression of the cohesin subunit Stromalin in the nervous system. The cohesin components Stromalin and NippedB are required for neuroblast death, and knockdown of Stromalin increases repressive histone modifications in neuroblasts. Thus Cut and cohesin regulate apoptosis in the developing nervous system by altering the chromatin landscape.

**Summary statement:** Cut regulates the programmed death of neural stem cells by altering cohesin levels and promoting a more open chromatin conformation to allow cell death gene expression.

## Introduction

Programmed cell death is important for normal nervous system development in organisms ranging from C. elegans to humans (Arya and White, 2015). Precise control of cell death in the nervous system requires the integration of spatial, temporal and cell identity signals from both cell intrinsic and extrinsic sources. Conserved signaling pathways that are instrumental in many developmental cell fate decisions also control the commitment of cells to death. Examining how these pathways interact to specify the cell death fate in a specific context is critical not only for understanding normal development but also to gain insight into how developmental pathways and homeostasis are disrupted in diseases such as cancer and neurodegeneration.

The RHG genes, *reaper* (*rpr*), *hid*, *grim* and *sickle* (*skl*) are required for virtually all cell death in the Drosophila embryo (White et al., 1994; Tan et al., 2011). These genes are transcriptionally activated in various combinations in cells fated to die. In the genome, the RHG genes are clustered in a 270kb death gene locus that is largely devoid of other genes. The large intergenic regions between the genes are highly conserved and contain cell type and temporally specific regulatory elements capable of activating different combinations of RHG genes to initiate cell death in specific developmental contexts (Bangs et al., 2000; Moon et al., 2008; Zhang et al., 2008; Tan et al., 2011; Arya and White, 2015).

To gain insight into the transcriptional regulation of cell death, we conducted a forward screen for genes required for the death of neural stem cells or neuroblasts (NBs) in the developing ventral nerve cord (VNC). A subset of NBs in the abdominal segments of the VNC is eliminated by apoptosis late in embryonic development (Truman and Bate, 1988; White et al., 1994; Peterson et al., 2002). In the absence of this death, the VNC becomes massively hypertrophic, and adult longevity is compromised (Peterson et al., 2002). We previously described how the Hox gene *abdominalA* (*abdA*) and *Notch* (*N*) are necessary and sufficient for NB death in the abdominal segments of the embryonic VNC (Arya et al., 2015). N activation in NBs requires the expression of the Delta ligand on NB progeny, and is required for a late pulse of *abdA* in NBs. *abdA* is necessary for abdominal NB death, and the late pulse of abdA could convey both spatial and temporal information about the specific NBs fated to die. Mis-expression of *abdA* is sufficient to cause ectopic NB death.

*abdA* regulates *rpr*, *grim* and *skl* expression through a regulatory element between *rpr* and *grim* called the Neuroblast Regulatory Region enhancer1 (enh1) (Arya et al., 2015). This element is required for the expression of *rpr*, *grim* and *skl* in NBs (Tan et al., 2011). Recent data indicate that *abdA*, *grainyhead* (*grh*) and *Su(H)*, downstream of *N* pathway activation, may be direct regulators of this neuroblast cell death enhancer (Khandelwal et al., 2017). However, it is clear that not all cells that express *abdA* and/or *grainyhead* activate the cell death genes and undergo cell death (KH and KW, unpublished observations). Thus, input from other factors must be required for the activation of NB death.

Here we report that the DNA binding protein Cut is required for NB death, acting through a mechanism distinct from AbdA and enh1. Cut is a transcriptional regulator with 4 DNA binding domains: 3 CUT domains and a Homeobox domain (Nepveu, 2001). Drosophila Cut is structurally and functionally homologous to Cux/CCAAT displacement protein (CDP) in human and Cux 1,2 in mouse, and can act as either an enhancer or repressor of transcription. In the Drosophila embryo, *cut* is expressed in the embryonic central and peripheral nervous system, Malpighian tubules and anterior and posterior spiracles (Blochlinger et al., 1990; Zhai et al., 2012). Loss of *cut* in the fly can enhance tumor growth, and *cut* has also been implicated in promoting differentiation and cell survival in posterior spiracle and tracheal development (Zhai et al., 2012; Pitsouli and Perrimon, 2013; Wong et al., 2014). In mammals, the functions of the Cux1 and Cux2 homologues are equally complex. Loss of Cux1 in mouse results in reduced proliferation and organ hypoplasia (Sansregret and Nepveu, 2008), but Cux1 has also been implicated as a haploinsufficient tumor suppressor in myeloid malignancies, and is associated with poor prognosis (Wong et al., 2014). Paralleling our findings on the role of Drosophila *cut* in NB death, Cux2 is required to limit the expansion of neuronal precursors in mouse brain development (Cubelos et al., 2008), but conversely in the spinal cord it is required for the maintenance of neural progenitors (Iulianella et al., 2008).

To examine how *cut* regulates cell death, we placed it in the regulatory framework defined by our previous studies (Arya et al., 2015). Our data indicate that *cut* plays a permissive role in neural stem cell apoptosis, acting to modify the chromatin landscape of the *rpr* region, to facilitate the expression of *rpr* and *grim* independently of the previously identified NB enhancer. We show that there is normally a temporal progression of NBs from an H3K27me3 low to an H3K27me3 high state, and H3K27me levels are enhanced throughout this progression in the absence of *cut*. In the cell death gene locus, this suppresses proapoptotic gene expression.

Importantly, we found that *cut* regulates expression of the cohesin subunit *stromalin* (*SA*). Cohesin is important for sister chromatid cohesion and long-range enhancer promoter interactions (Rollins et al., 1999; Kagey et al., 2010). We demonstrate that cohesin components are required for normal NB death, and that loss of cohesin results in increased numbers of NBs with high levels of H3K27me3. We propose a model for the regulation of NB death through the combinatorial control of chromatin accessibility, chromatin architecture, and the temporal and spatial activity of sequence-specific transcription factors.

## Results

### *cut* is necessary and sufficient for abdominal NB death

The *cut* gene was identified in an RNAi screen for regulators of NB death (Arya et al., 2015). Knockdown of *cut* in the CNS with multiple RNAi lines results in a large increase of persistent NBs late in Drosophila embryogenesis (Fig. 1B), at a time when the majority of abdominal NBs have undergone apoptosis in the wild type (Fig. 1A). Ectopic abdominal NB survival is also detected in 3^rd^ instar larvae (Fig. S1). Embryos homozygous for a *cut* null mutant, *cut*^*C145*^ (Johnson and Judd, 1979; Micchelli et al., 1997) also show persistent abdominal neuroblasts in late embryogenesis (Fig. 1C). The rescue of NB death is not due to *cut* activity in neighboring glia, as *cut* knockdown in glia does not inhibit NB death (Fig. S1). Thus, *cut* is required for the programmed death of NBs in late embryogenesis.

**Figure 1.**
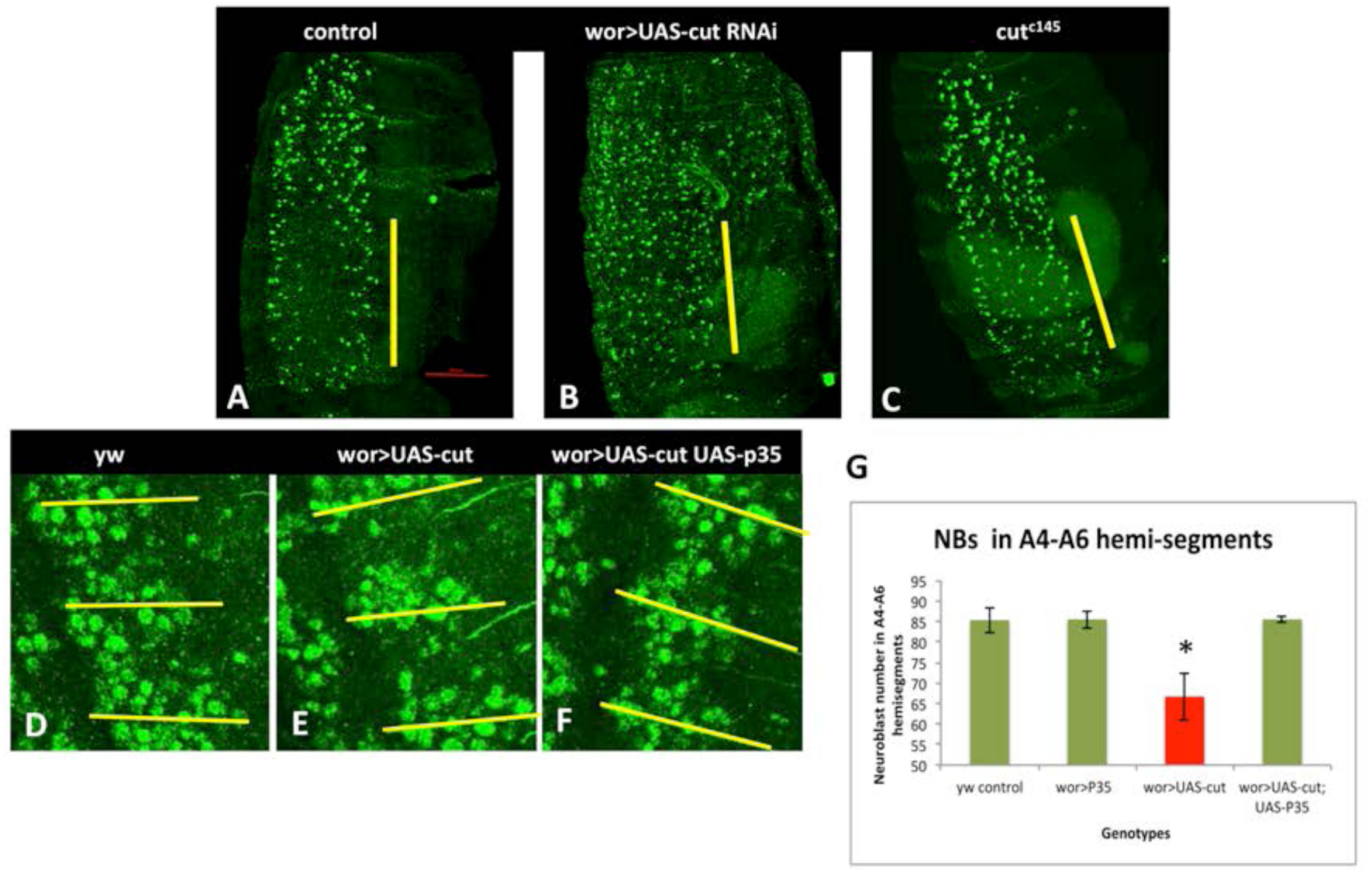
*cut* is necessary and sufficient for NB death. **A-C**) Knockdown of *cut* in the nervous system, or *cut* loss in mutants embryos, results in ectopic NB survival, as detected by Dpn staining in stage 17 embryos. D-F) *cut* overexpression results in NB loss by apoptosis. Fewer NBs can be seen in each hemisegment, particularly in the anterior of each hemisegment. This NB loss is blocked by the broad-spectrum caspase inhibitor p35. G) Premature NB loss resulting from *cut* overexpression is significant at stage 14.* P<0.05 by unpaired T test

*cut* is expressed in the CNS starting at early stage 12. Initial expression is very low, and is strongly expressed in many cells of the CNS, including NBs, by stage 15 (Fig. S2). Thus *cut* is expressed in NBs at a time when NB death begins (stage 14), and is expressed in most or all NBs at the time cell death peaks. Widespread *cut* expression in both NBs and neurons indicates that at normal levels, *cut* is not sufficient to activate apoptosis in all cells. Rather, *cut* may be permissive for the activation of the cell death genes by additional spatial and temporal factors, including *N*, *abdA* and *grh* (Cenci and Gould, 2005; Maurange et al., 2008; Arya et al., 2015; Khandelwal et al., 2017).

Overexpression of *cut* in the CNS results in premature loss of NBs in abdominal segments (Fig. 1E, G). We see loss of many abdominal NBs at stage 14, before they normally die. This *cut*-induced NB loss can be inhibited by the baculovirus broad-spectrum caspase inhibitor p35, demonstrating that NB loss is due to caspase-dependent cell death, and not to alterations in NB fate (Fig. 1F, G). Overexpression of *cut* in the whole embryo with heatshock-gal4 also results in ectopic cell death (Fig. S3). Taken together, these findings demonstrate that *cut* is necessary for timely NB death, and is temporally limiting for the activation of cell death.

### *cut* acts upstream of rpr and grim and downstream of enh1

NB death requires the activity of the *rpr*, *grim* and *skl* genes (Tan et al., 2011). These genes are transcribed in doomed cells, and the Rpr, Grim and Skl proteins inhibit DIAP1 to activate caspases (Kornbluth and White, 2005). To determine whether *cut* regulates NB death through this pathway, we assessed *rpr* and *grim* transcript levels in the absence of *cut* and when *cut* is overexpressed. In stage 15 embryos both *rpr* and *grim* expression are clearly reduced when *cut* is knocked down in the CNS (Fig. 2A-D). Conversely, *cut* overexpression throughout the CNS with wor-gal4 results in a substantial increase in levels of *rpr* and *grim* transcripts when cell death is inhibited with p35 (Fig. 2E-H). Interestingly, even when *cut* is overexpressed in many or all cells of the CNS, *rpr* and *grim* are hyper-activated only in a subset. This suggests that *cut* is permissive for *rpr* and *grim* expression, but requires additional regulators to fully activate the NB death program.

**Figure 2.**
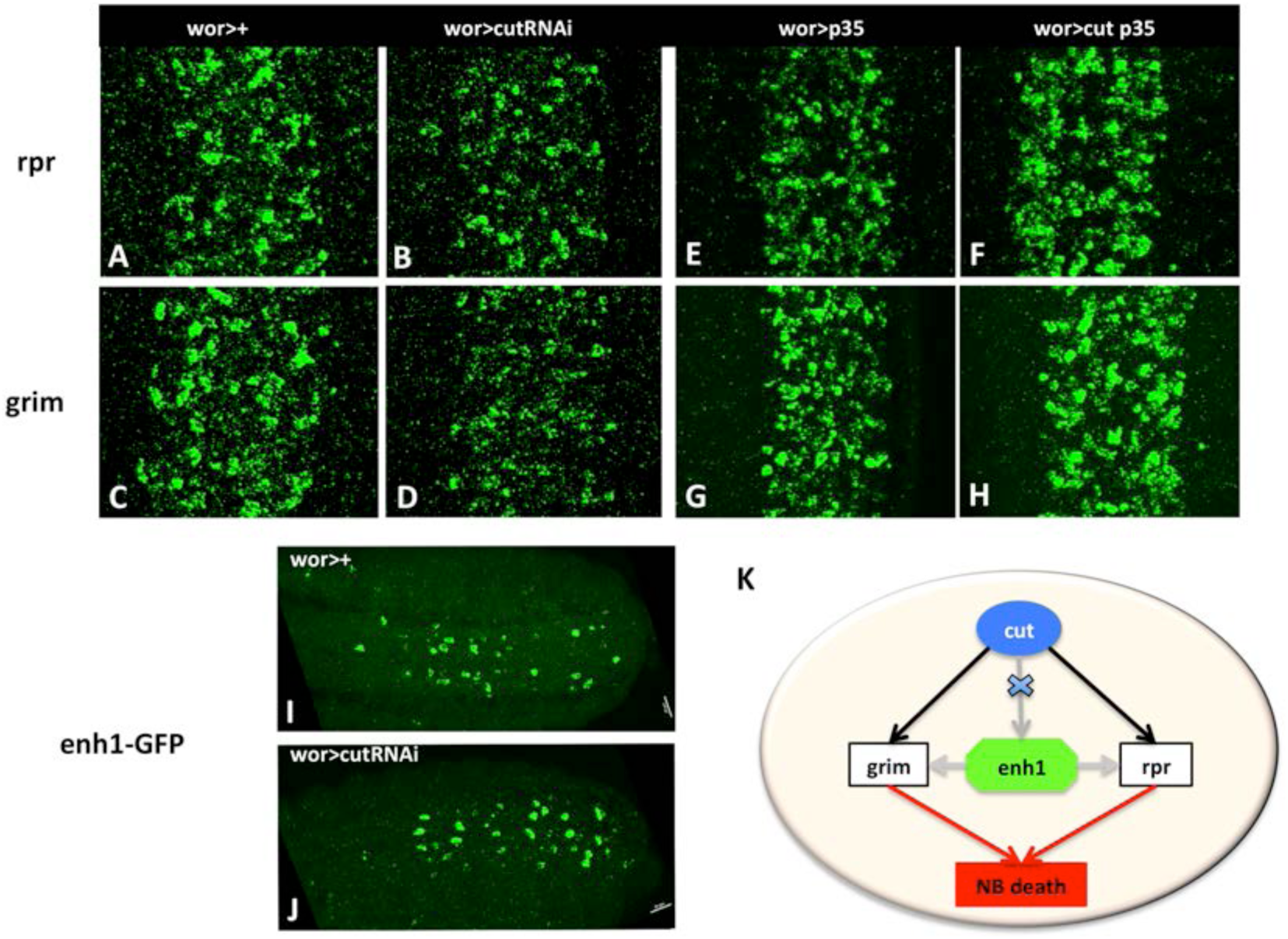
*cut* alters rpr and grim levels independently of the neuroblast regulatory region. A-D) *cut* knockdown in the CNS decreases *rpr* and *grim* expression, as detected by in situ. E-H) On *cut* overexpression, *rpr* and *grim* mRNA levels are increased. P35 is used to block ectopic cell death induced by *cut*. I-J) *cut* knockdown does not alter expression of enhancer1-GFP, indicating that *cut* is likely to influence *rpr* and *grim* expression and cell death independently of the NB regulatory region (K).

Our previous studies identified a regulatory region between *rpr* and *grim*, the Neuroblast Regulatory Region, which controls *rpr*, *grim* and *skl* expression to promote abdominal NB death (Tan et al., 2011). A 5kb transgenic reporter generated from this region, enh1-GFP, is expressed in doomed abdominal NBs and is responsive to the levels of *N* and *abdA*, which regulate NB death (Arya et al., 2015). We found that *cut* does not regulate enh1-GFP activity: knockdown of *cut* does not decrease enh1-GFP expression. Instead, we see an increase in the number of enh1 expressing cells on *cut* knockdown (Fig. 2 I, J). This suggests that loss of *cut* blocks the death of enh1-GFP expressing cells. Furthermore, *cut* overexpression does not increase enh1-GFP levels, and is able to induce NB death in embryos that lack the NB enhancer1 due to the MM3 deletion (Tan et al., 2011) (data not shown). Therefore, *cut* acts independently of enh1 to facilitate the activation of *rpr* and *grim* for NB death (Fig. 2K).

### *cut* acts downstream of *abdominalA*

*abdA* is necessary and sufficient for abdominal NB apoptosis (Prokop et al., 1998; Arya et al., 2015; Khandelwal et al., 2017). Overexpression of *abdA* results in ectopic NB death and enhanced and ectopic enh1-GFP expression. If *cut* acts to regulate apoptosis downstream of *abdA*, *cut* knockdown should block killing by ectopic *abdA*. Indeed, we found that *cut* knockdown in the context of *abdA* overexpression blocks ectopic NB death in both thoracic and abdominal segments of the VNC (Fig 3A-C). Importantly, ectopic *abdA*-activated enh1-GFP expression is still apparent in *cut* knockdown (Fig. 3D-F), indicating that *cut* does not prevent *abdA* from activating the regulatory region, but blocks *rpr* and *grim* activation by the enhancer. We also found that *abdA* knockdown does not rescue NB death induced by *cut* overexpression (Fig. 3G-J). These data support the conclusion that *cut* regulates NB death downstream of *abdA* and enh1, and upstream of the RHG genes.

**Figure 3.**
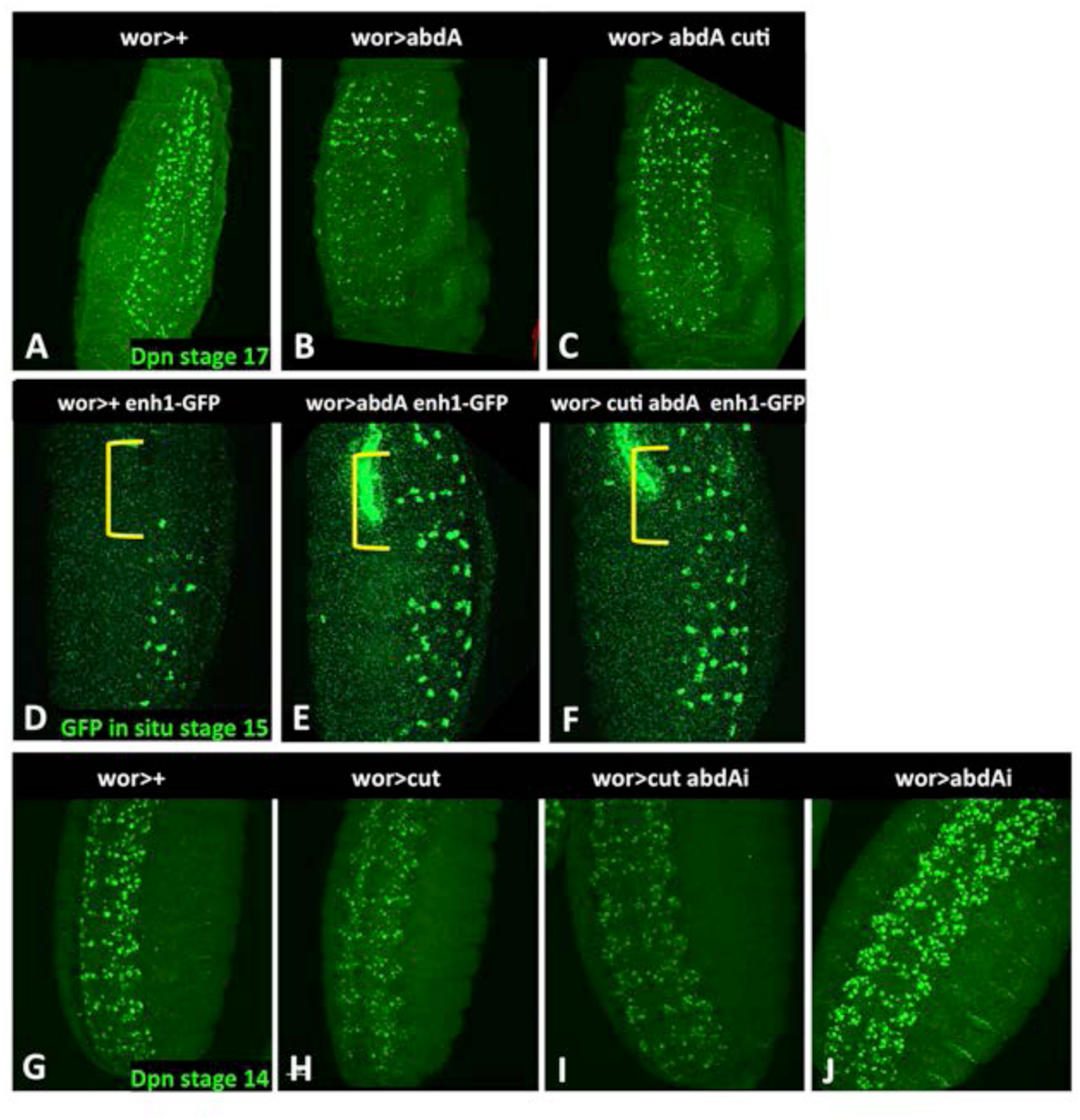
*cut* acts downstream of abdA. A-C) Knockdown of *cut* inhibits NB killing by *abdA* overexpression in both abdominal and thoracic domains. NBs are detected by anti-Dpn. D-F) Loss of *cut* does not inhibit ectopic enh1 expression in thoracic segments (bracket) induced by *abdA* mis-expression. GFP is detected by in situ, G-H) Knockdown of *abdA* does not rescue NB death induced by *cut* overexpression. *abdA* knockdown alone results in ectopic NB survival (J).

### *cut* does not inhibit NB death through a binding site in the IRER left barrier

Enhancer/promoter interactions can be temporally controlled by changes in chromatin accessibility (Uyehara et al., 2017). Loss of *cut* inhibits NB death downstream of enh1 activity, suggesting that *cut* could influence *rpr* and *grim* expression in NBs by influencing chromatin accessibility in the *rpr*/*grim* region. Another enhancer in the death gene locus, the irradiation responsive enhancer region (IRER), which is located 5’ of the *rpr* promoter, shows temporal changes in chromatin conformation that are responsible for the reduced sensitivity to irradiation in later stages of embryogenesis (Zhang et al., 2008). Previous studies showed that in older embryos there is an abrupt change in chromatin conformation at the promoter proximal end of the IRER, with a sharp decrease in H3K27me3 and H3K9me3 and an enrichment of H3K4me3 at the *rpr* proximal promoter (Fig. S4) (Lin et al., 2011). This finding is consistent with our hypothesis that changes in chromatin conformation at the death gene locus regulate competence to respond to apoptosis-inducing signals, but does not address the biological role of the IRER in the regulation of NB death.

The previous study identified a chromatin barrier within the IRER (IRER left barrier element, or ILB) containing a putative Cut binding site that was necessary for barrier function. We therefore asked whether the Cut binding site in the ILB was critical to prevent heterochromatin spreading from the IRER into the *rpr* proximal promoter, therefore allowing *rpr* activation and NB death in response to activation of enh1. We generated several deletions of the putative Cut binding site using CRISPR/Cas9 (Fig. S4). We examined NB death in animals homozygous for these deletions, and found that NB death was normal (Fig. S4). NB death is also normal in animals homozygous for the larger IRER deletion generated by the Zhou lab (data not shown). Thus the predicted Cut binding site in the ILB is not required for NB death downstream of enh1 activation. This result suggests that additional *cut*-dependent mechanisms could mediate communication between the NB enhancer and the *rpr* and *grim* promoters.

### *cut* inhibits NB death by altering repressive chromatin in NBs

We hypothesized that an increase in repressive histone modifications at the death gene locus in NBs could limit NB death on *cut* knockdown. To test this, we assayed repressive and activating histone modifications by ChIP in control embryos and after CNS-specific *cut* knockdown. To enrich for NB chromatin and to limit stress-induced changes in cell death gene expression, ChIP was carried out on chromatin isolated from sorted fixed CNS nuclei from wor>dsRed (wor>+) and wor>dsRed cutRNAi (wor>cutRNAi) embryos (Bowman et al., 2013).

In contrast to data previously obtained from whole stage 16 embryos (14-16 h) (Negre et al., 2011), we noted that the overall enrichment for H3K27me3 was generally low in the *grim* to *rpr* region in CNS chromatin. In response to *cut* knockdown, we saw a slight enrichment for this repressive mark in this region (Fig. 4A), consistent with the decreased transcription of *grim* and *rpr* we detected in the absence of *cut* (Fig. 2). This suggests that *cut* may normally inhibit the formation of facultative heterochromatin in the *grim* to *rpr* region in the developing CNS. No major alterations in H3K27me3 were detected in the *bithorax* complex in any of our experiments (data not shown), indicating that *cut* does not regulate H3K27me3 levels at all genes in the CNS.

**Figure 4.**
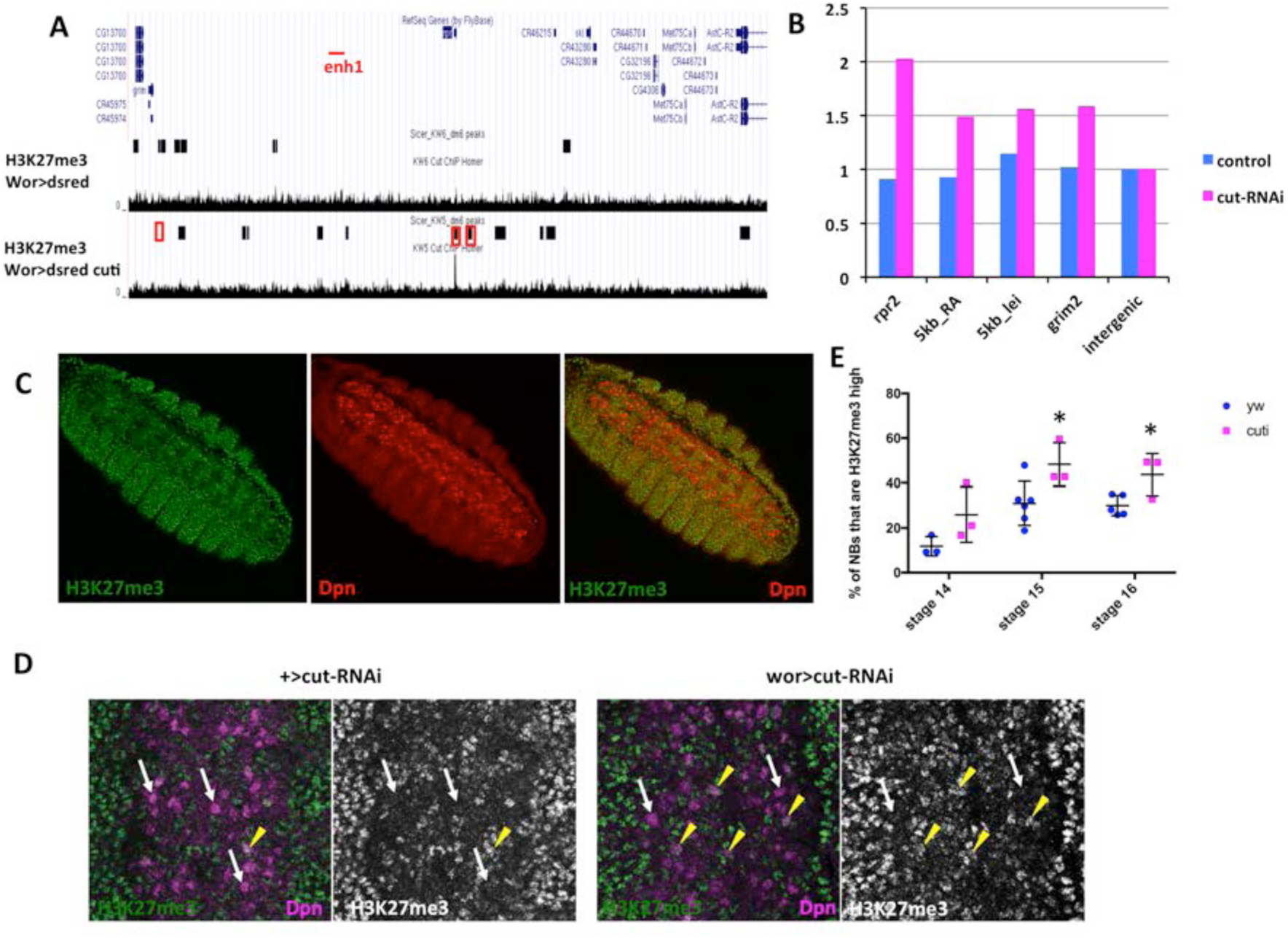
*cut* knockdown increases H3K27me3 levels in the rpr to grim interval. A) ChIP-Seq on sorted CNS nuclei from wor>+ and wor>cutRNAi show an increase in H3K27me3 modifications in the *rpr* region after *cut* knockdown, as detected by SICER peak calling. B) Verification by ChIP-qPCR shows increased enrichment of H3K27me3 at the promoters of *grim* and *rpr*, and 5’ of *rpr* in chromatin from sorted nuclei isolated from wor>cutRNAi when compared to wor>+. All data are normalized to levels in the intergenic region. Primer regions are boxed in A. C) H3K27me3 levels are lower in NBs than in the rest of the embryo. D,E) The proportion of NBs with strong H3K27me labeling show increases at embryos age (white arrows, H3K27me3-negative NBs; yellow arrowheads, H3K27me3-high NBs). Knockdown of *cut* increases the number of H3K27me3 high NBs at stages 14 through 16. * p<0.05 by unpaired T test.

To confirm changes detected by ChIP-seq, we assayed H3K27me3 enrichment at several positions in the *rpr* region by ChIP-qPCR on independent chromatin preparations from fixed CNS nuclei. We repeatedly found that *cut* knockdown led to enrichment for H3K27me3 within the *rpr* and *grim* open reading frames, in the region 5kb upstream of *rpr* (Fig 4B), and at the *rpr* promoter (Fig. S5). We conclude that loss of *cut* increases repressed histone modifications in the RHG region in the developing CNS, and this could be responsible for decreased *rpr* and *grim* expression. However, the effect of *cut* knockdown is relatively limited, which could reflect redundant mechanisms underlying the activity of *cut*, or could be due to the low representation of the cells of interest (NBs) in our chromatin preparation.

To focus more precisely on *cut*-dependent changes in histone modifications in NBs, we stained control and *cut* knockdown embryos for H3K27me3. Strikingly, we found that in control embryos at stage 14, there was a clear difference in the overall levels of H3K27me3 in NBs, as compared to other tissues in the embryo, and to other cells in the CNS (Fig 4C). H3K27me3 levels were low or undetectable in the ventral NB layer, and much higher in the more dorsal layers of the CNS containing the differentiated neurons and glia. This is not due to a defect in histone antibody accessibility in these cells, as other histone modifications were not strikingly different in NBs compared to other neural cells (Fig. S6). Quantification of NBs with high H3K27me3 showed that only 12% of NBs at stage 14 had high levels of H3K27me3 (Fig. 4D-E). The lower levels of the repressive H3K27me3 modification in early NBs may reflect the increased plasticity of chromatin in these stem cells (Marshall and Brand, 2017).

Furthermore, we found that levels of H3K27me3 increase in control NBs over time, so that by stage 16 about 30% of NBs were scored as H3K27me3 high. This increase in repressive histone modifications could reflect a restriction of stem cell potential over developmental time. At all stages, the lower levels of H3K27me3 in NBs as detected by staining could explain the relatively low levels of H3K27me3 peaks detected in our ChIP experiments on CNS chromatin.

Surprisingly, we found that *cut* knockdown increased the number of NBs with high H3K27me3 at stages 14 through 16 (Fig. 4D-E). This indicates that *cut* in NBs is required to hold chromatin in a more open conformation by inhibiting the deposition of H3K27me3. This open conformation could be required for normal *rpr* and *grim* activation during the period of cell death. Loss of *cut* did not prevent the temporal increase in H3K27me3 levels, as NBs in older *cut* knockdown embryos had a higher number of H3K27me3-high NBs than earlier stages.

### Cohesins operate downstream of *cut* to regulate NB death

Because *cut* has not been characterized as a histone modifier or as part of a histone modifier complex, we hypothesized that the effect of *cut* knockdown on H3K27me3 levels in NBs was likely to be indirect. We scanned our H3K27ac ChIP-seq data to identify potential *cut*-regulated genes that could restrain H3K27me3 levels in NBs. Most genes did not exhibit changes in H3K27ac peaks, including Polycomb complex components. However, two structural components of the cohesin complex, *stromalin* (*SA*) and *SMC1* showed decreased H3K27ac levels following *cut* knockdown (Fig. 5A). The cohesin complex is implicated in three-dimensional chromatin architecture, including enhancer-promoter interactions (Kagey et al., 2010). Cohesin also interacts with the Polycomb Repressive Complex 1 (PRC1), and may sequester PRC1 from repressive chromatin, resulting in an overall mutually exclusive distribution of cohesin binding and repressive histone modifications (Misulovin et al., 2008; Schaaf et al., 2013).

**Figure 5.**
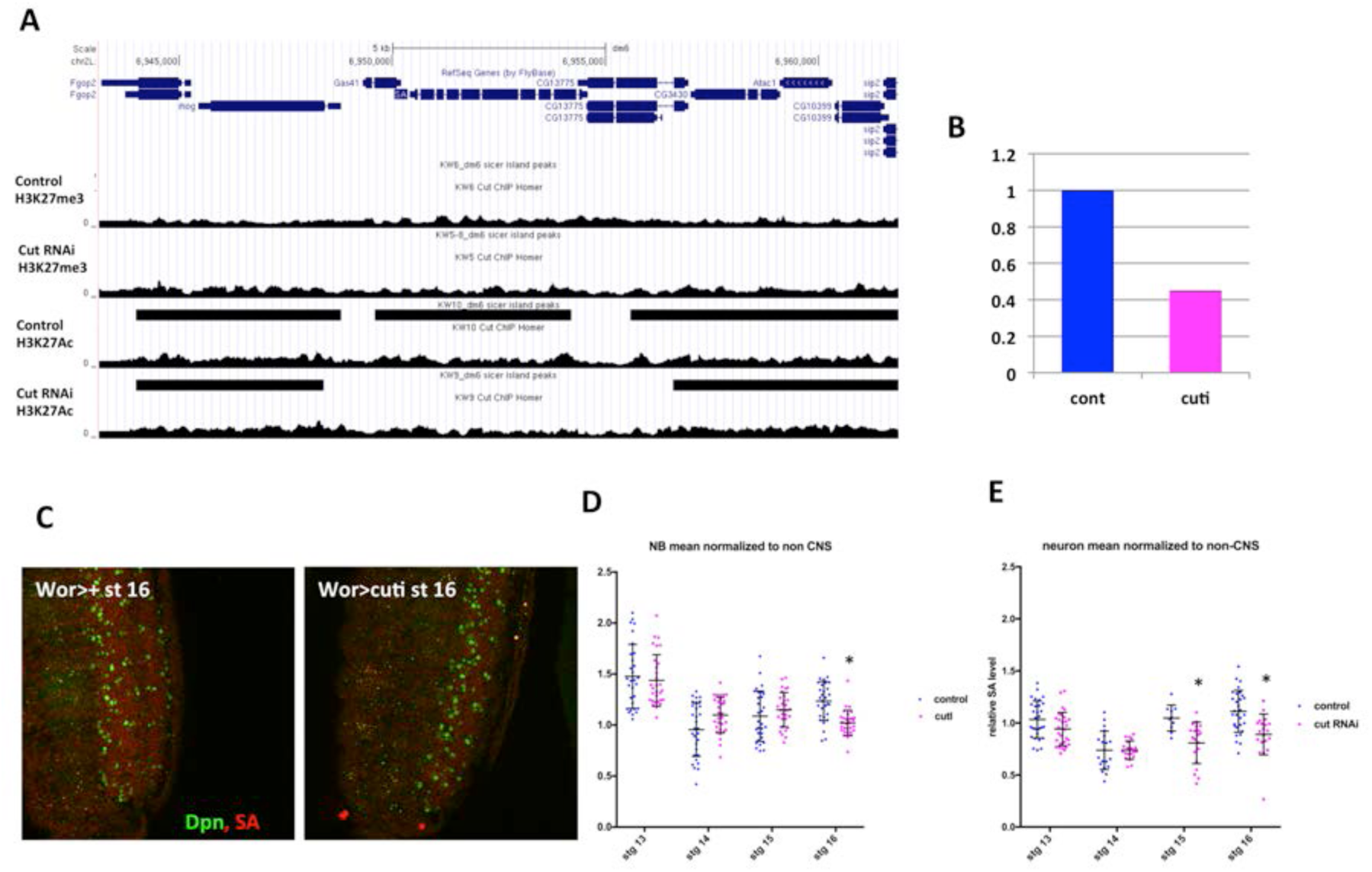
*cut* knockdown alters the expression of the cohesin component SA. A) A peak of H3K27Ac over *SA* disappears on *cut* knockdown. B) qPCR on RNA from sorted CNS nuclei shows a decrease in *SA* levels. C) SA levels are decreased in the CNS by *cut* knockdown. D) mean intensity of SA in NBs relative to non-CNS cells is significantly decreased on *cut* knockdown. E) mean intensity of SA in neurons relative to non-CNS cells is significantly decreased on *cut* knockdown. * p<0.05 by unpaired T test.

We hypothesized that *cut* knockdown could decrease cohesin activity, leading to increased repressive chromatin and decreased cell death gene expression. A decrease in cohesin expression upon *cut* knockdown could interfere with communication between the NB enhancer and the *rpr* and *grim* promoters, altering their expression and inhibiting NB death. In addition, loss of cohesin could enhance the deposition of repressive chromatin to limit the expression of cell death genes.

To examine whether *SA* expression was controlled by *cut*, we assayed *SA* RNA levels by qPCR on RNA prepared from sorted CNS nuclei from control and wor>cutRNAi embryos (Fig. 5B). *SA* RNA levels were decreased on *cut* knockdown (Fig. 5B). Furthermore, decreased *SA* protein levels were detected on *cut* knockdown in both NBs and neurons (Fig. 5C-E). In contrast, Cut protein levels in the CNS were not altered by *SA* knockdown (Fig. S7).

If *cut* regulates NB death by altering cohesin expression, then cohesin knockdown should phenocopy loss of *cut* and inhibit NB death. Indeed, we found that knockdown of *SA* results in ectopic NB survival in late embryos (Fig. 6A, B, D). In addition, knockdown of *NippedB*, part of the kollerin complex required for cohesin loading (Dorsett and Kassis, 2014), resulted in ectopic NB survival (Fig. 6C). These data demonstrate a previously unknown requirement for cohesin in the regulation of NB death.

**Figure 6.**
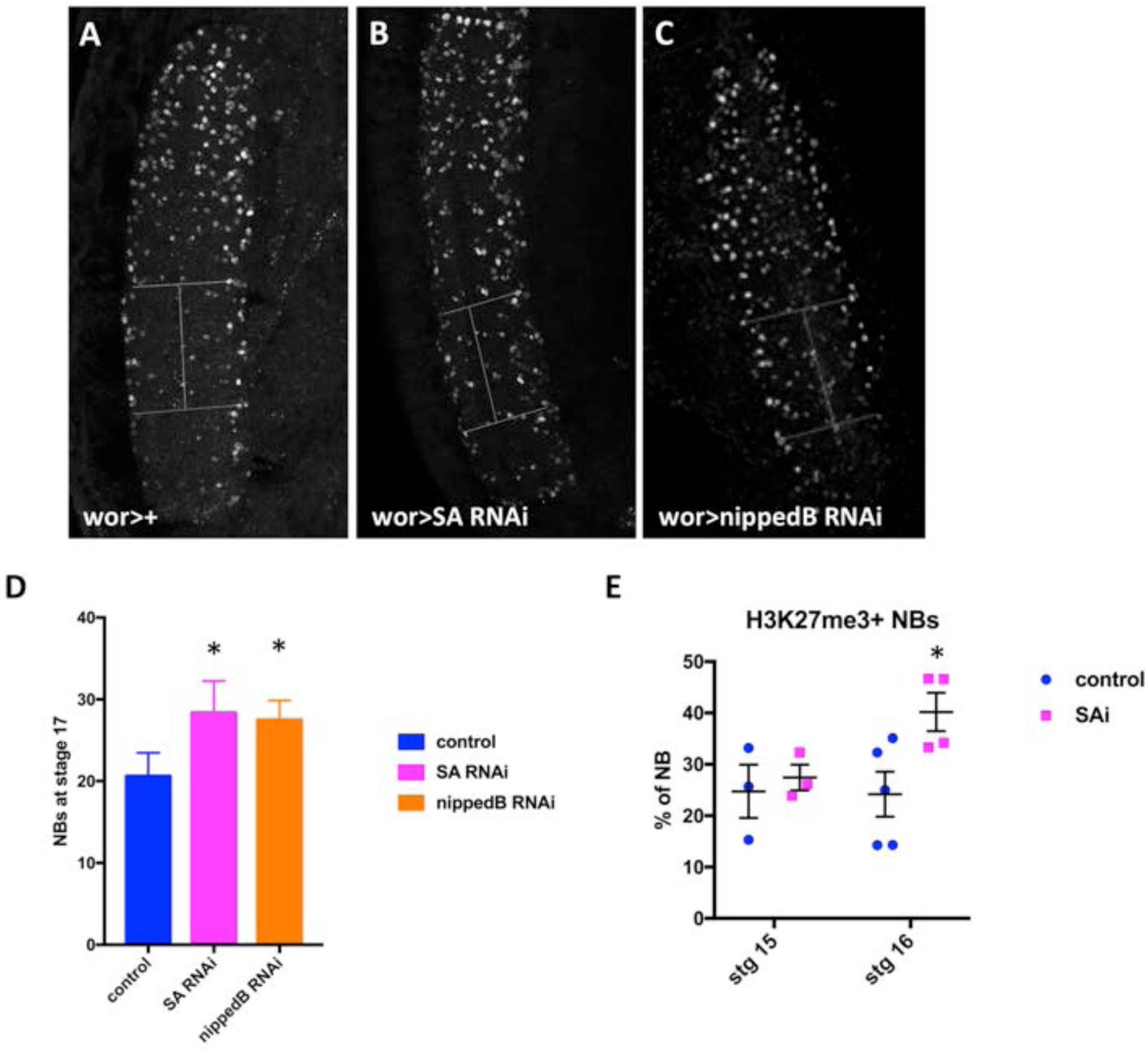
Cohesin is required for normal abdominal NB death. A-C) Dpn staining reveals ectopic NB survival in stage 17 embryos after *SA* or Nipped-B knockdown. D) Dpn positive NBs were counted in 3 segments, and show significant increases in stage 17 embryos on cohesin knockdown. E) Knockdown of *SA* results in an increased proportion of H3K27me3 positive NBs at stage 16. *p< 0.05 by unpaired T test.

Cohesin knockdown increases Pc binding at the majority of H3K27me3 marked genes (Schaaf et al., 2013). We examined overall H3K27me3 levels in NBs after cohesin knock down. We found that *SA* knockdown increased the number of NBs with high levels of H3K27me3 in embryos (Fig. 6E), suggesting that higher H3K27me3 levels in NBs on *cut* knockdown could be caused by decreased cohesin. Thus, cohesin is required for abdominal NB death, and may regulate cell death gene expression by altering overall levels of repressive chromatin in NBs (Fig. 7).

**Figure 7.**
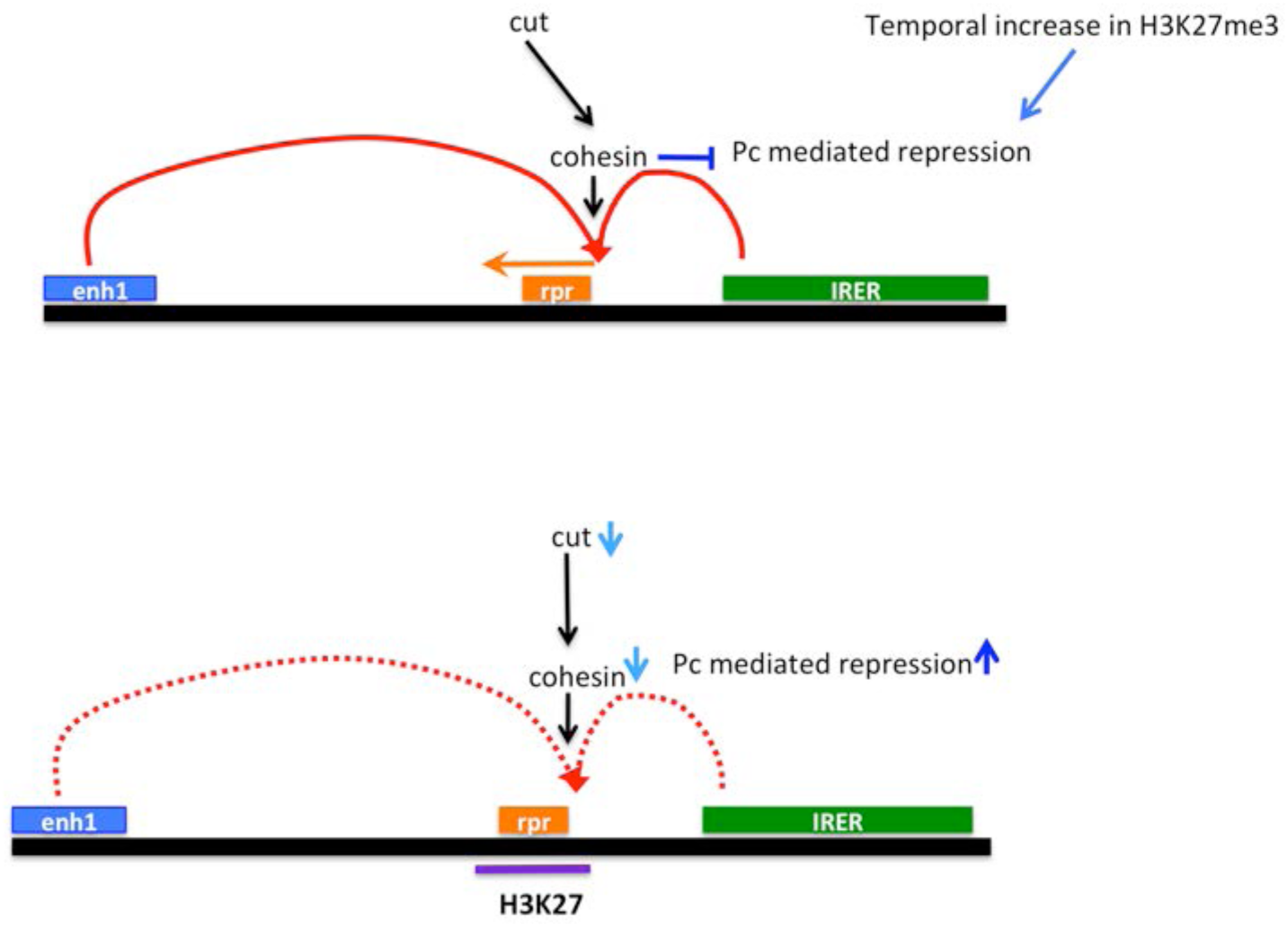
Model of *cut* action in NB death. As neural stem cells age they show an overall increase in repressive chromatin, marked by H3K27me3. Expression of *cut* inhibits this increase, at least in part through enhancement of cohesin expression. At the *rpr* promoter, this allows enhancers, activated by additional cell type specific spatial and temporal factors, to turn on the transcription of *rpr*, *grim* and *skl*. When *cut* expression is suppressed, *SA* and possibly other cohesin subunits decline, allowing a premature increase in H3K27me3 in NBs. At the *rpr* locus, this blocks expression in response to upstream apoptosis regulatory factors. Other upstream regulators may influence cohesin levels in other tissues.

## Discussion

In this work we report that *cut* plays a previously unknown role in the regulation of NB death. *cut* is permissive for the expression of *rpr* and *grim*, acting downstream of the previously identified neuroblast regulatory region. We show that *cut* loss increases the number of NBs with high levels of H3K27me3, indicating a role for *cut* in maintaining open chromatin in NBs. At the RHG locus, this is reflected in higher levels of H3K27me3, associated with lower *rpr* and *grim* expression. Importantly, we find that *cut* regulates the levels of the cohesin subunit *SA* in the CNS, and we show that cohesin is required for NB death. This work demonstrates a novel connection between *cut* and cohesin in controlling the chromatin landscape and cell death in the developing CNS.

### Is *cut* the “cell identity” signal that is permissive for NB death?

Our previous work identified the Hox gene *abdA* as an important spatial signal for NB death in the embryo (Arya et al., 2015). A late pulse of AbdA in NBs is regulated by *Notch* activation that is dependent on Delta ligand expression in NB progeny. *abdA* is necessary and sufficient for NB death, and has been shown to bind to enh1 in the Neuroblast regulatory region (Khandelwal et al., 2017). However, *abdA* is clearly expressed in many cells that do not die (Karch et al., 1990; Arya et al., 2015). Furthermore, mis-expression of *abdA* does not activate ectopic NB death prior to stage 13 of embryogenesis (Prokop et al., 1998; Arya et al., 2015), suggesting that there are temporal and cell identity signals that regulate the competence of cells to respond to *abdA*.

Here we identify *cut* as a novel regulator of NB death. Expression of *cut* in the embryonic CNS increases as NB death begins. However, most cells that normally express *cut* do not die, indicating that other factors coordinate with *cut* to regulate NB death. We find that loss of *cut* inhibits *rpr* and *grim* transcription, but in contrast to *abdA* and N, *cut* does not act on enh1, as detected by enh1-GFP. In addition, *cut* knockdown blocks NB killing in response to *abdA* mis-expression, despite an expansion of enh1 expression. These data indicate that *cut* acts downstream of enh1, and suggests that *cut* acts in the nervous system as a permissive factor that regulates the competence of NBs to respond to other cell death signals.

### *cut* alters the chromatin landscape in the nervous system

We found that *cut* functions in the CNS to restrict overall levels of repressive H3K27me3-marked chromatin. We demonstrate that NBs have a significantly lower level of overall H3K27me3 than other tissues in the embryo, possibly associated with stem cell plasticity (Zhu et al., 2013). As embryos age, the number of NBs with high overall levels of H3K27me3 increases. The cause and consequences of this transition are unknown, but could be related to a gradual restriction of NB fate (Yuzyuk et al., 2009; Zhu et al., 2013; Marshall and Brand, 2017).

We found that loss of *cut* promotes more NBs to acquire an H3K27me3 high state throughout later stages of embryogenesis. Interestingly, in both control and *cut* knockdown there is a temporal increase in the proportion of NBs with high H3K27me3. This suggests that additional temporal factors control this maturation of NBs to a more repressed state, but *cut* restrains the number of H3K27me3 high cells throughout this transition.

Our data indicate that *cut* overexpression is sufficient to cause increased *rpr* and *grim* expression and apoptosis in NBs. This is not due to hyper-activation of enh1, as ectopic *cut* does not increase enh1-GFP expression and can cause NB death even in the absence of the neuroblast regulatory region. In addition, *cut* overexpression causes increased cell death in other cells that normally survive, as seen with heat shock-gal4. This suggests that ectopic *cut* could activate additional upstream apoptosis-inducing signals, directly activate *rpr* and *grim* expression, or could open the *rpr* region for activation by regulators that do not normally activate *rpr* and *grim*.

The role of *cut* in activating NB death is in contrast to previous work suggesting that *cut* inhibits cell death in the developing posterior spiracle by directly inhibiting *rpr* expression (Zhai et al., 2012). In the developing spiracle, *cut* is also required for normal differentiation. Several other tissues also require *cut* for normal differentiation, such as the bristle cells in the eye, and the developing trachea. In these tissues, cell death is also increased in the absence of *cut* (Pitsouli and Perrimon, 2010; Zhai et al., 2012). The role of *cut* in promoting cell survival in these tissues differs from its role in facilitating cell death in the CNS. This may reflect the diverse activities of *cut* as a transcriptional regulator, or could be due to *cut*’s activity as a chromatin organizer, altering the landscape for binding by both activators and repressors of RHG gene transcription. Both pro-differentiation and pro-apoptotic roles of *cut* are consistent with its role as a potential tumor suppressor (Zhai et al., 2012; Wong et al., 2014).

### Cell death genes are highly sensitive to altered chromatin accessibility

This study, and previous work from the Zhou lab, indicates that the *rpr* region is particularly sensitive to alterations in chromatin conformation, reflecting the need for rapid and robust transcription of the cell death genes in cells fated to die. Other factors that control histone modifications are involved in cell death. For example the dUTX H3K27me3 demethylase is required for Ecdysone Receptor-mediated activation of *rpr* expression in salivary gland death (Denton et al., 2013). This supports our finding that a more open chromatin conformation is particularly important for cell death gene activation. Expression of other components of the cell death pathway may also be controlled by changes in chromatin conformation. For example, treatment of Drosophila larvae with HDAC inhibitors, or HDAC1 knockdown, increases sensitivity to cell death activation through altered expression of caspases (Kang et al., 2017). Conversely, loss of Polycomb-mediated suppression is associated with loss of postembryonic NBs, although this may be due to ectopic *abdA* expression (Bello et al., 2007). There is also evidence for epigenetic regulation of genes important for cell death in the mammalian nervous system and in cancer (Wright et al., 2007; Song et al., 2011). Here we provide evidence that control of histone modifications in the *rpr* region is an important aspect of developmental cell death regulation.

### Cohesin as a regulator of cell death

Given the lack of evidence for a direct histone-modifying role of Cut in regulating cell death, we investigated alternative indirect mechanisms and determined that *cut* promotes expression of the cohesin subunit *SA*. We found that, similar to loss of *cut*, down-regulation of *SA* or Nipped-B results in ectopic NB survival. Cohesins are involved in sister chromatin cohesion, formation of topologically associated domains and in long-range enhancer promoter interactions (Kagey et al., 2010; Newkirk et al., 2017). This latter function may be particularly important in Drosophila developmental cell death. Multiple cell death genes must be activated in different tissues in response to overlapping signals impinging on distinct regulatory enhancers (Jiang et al., 2000; Lohmann et al., 2002; Zhang et al., 2008; Arya et al., 2015; Khandelwal et al., 2017). This suggests that three dimensional chromatin interactions, including those mediated by cohesin, are critical for facilitating precise gene activation in the RHG region.

Loss of one copy of the human Nipped-B homolog NIBPL, and of other cohesin components, is associated with Cornelia de Lange syndrome, a developmental disorder affecting growth, cognitive function and facial and limb morphology (Wu et al., 2015; Newkirk et al., 2017). This is likely due to the downregulation of developmentally important genes, as detected in NIBPL +/-MEFs (Newkirk’17). Nipped-B heterozygous flies also exhibit reduced growth, learning and memory deficits, abnormal brain morphology and reduced expression of many genes (Wu et al., 2015). Interestingly, Nipped-B heterozygotes are resistant to dMyc induced apoptosis, a phenotype also seen in the IRER mutants (Wu et al., 2015; Zhang et al., 2015), suggesting that cohesin may also regulate cell death activated by the IRER enhancer. Our data suggest that control of cell death in the nervous system could also contribute to the Cornelia de Lange syndrome phenotype. Additional studies are needed to understand how cohesin activity is directed towards regulating the expression of specific genes.

Precise control of apoptotic gene expression is particularly important in the nervous system, the site of the majority of developmental cell death in flies, worms and mammals, and the tissue most affected by the absence of cell death (Arya and White, 2015). Our work has led to a greater understanding of the temporal, spatial and tissue specific control of this death in flies through developmentally important transcription factors as well as regulation of chromatin accessibility and architecture. Given the conserved function of the pathways we have identified, it is likely that these studies will provide insight into the regulation of cell death in human nervous system development and disease.

## Materials and Methods

### Embryo collection and nuclei preparation

Embryos were collected for 16 hr. at 25°. Dechorionated embryos were fixed in 1:1 solution of 1.8% formaldehyde and heptane for 15 min at room temperature. The fixative was quenched with a 2 min wash with 125 mM glycine in PBS with 0.1% Triton-X100 (PBS:130 mM NaCl, 7 mM Na2HPO4, 3 mM KH2PO4,, pH 8.0), and then briefly rinsed with PBS 0.1% Triton-X100. Embryos were snap frozen in liquid nitrogen, and stored at −80°C. About 1g of embryos of each genotype were used for nuclei isolation as described in Bowman et al. (Bowman et al., 2013; Bowman et al., 2014).

### Nuclear sorting and ChIP-seq

Fixed nuclei from wor>dsRed embryos were enriched using a Bio-RAD S3e cell sorter with 561nm excitation. Nuclei were sorted at 4^0^C in 100ul of PBS, with 1% BSA, 0.1% Triton-X and 1X protease inhibitor. wor-gal4 is expressed in the nervous system from stage 11 onwards (Arya et al., 2015). The sorted nuclei represent approximately 0.5-3% of total embryonic nuclei, and were at least 50% pure, based on post-isolation assessment of ds-red by confocal microscopy. About a million nuclei were used for chromatin preparation. After isolation chromatin was fragmented with 15U of micrococcal nuclease (MNase, Worthington Biochemical) followed by 3 min sonication in a Diagenode Bioruptor 377 (Bowman et al., 2013). Immunoprecipitation was carried out with 2ug of H3K27me3 antibody (Active Motif 39136) or 1ug of H3K27Ac antibody (Active Motif, 39136). Single end tag libraries were prepared and sequenced on an Illumina, Hiseq2500 in high output mode at the MGH Next Generation Sequencing Core).

### ChIP-seq data analysis

High throughput sequence data were processed and analyzed for quality. Samples with reasonable ChIP strength were further analyzed. Reads were mapped to the genome (dm6) with Bowtie2 (Langmead and Salzberg, 2012). The resulting SAM files were used to identify enrichment using MACS2 (Feng et al., 2012) and SICER (Xu et al., 2014). The resulting BED file of enriched genomic regions and the normalized BedGraph files were loaded to the UCSC Genome Browser for comparison and analysis.

### ChIP-qRT-PCR analysis

For the validation of ChIP-Seq data, ChIP-qPCR was performed. Immunoprecipitation conducted as described above using the H3K27me3 antibody. The immunoprecipitated DNA was processed for qPCR analysis using iTaq universal SYBER green supermix (Bio-Rad, CA) on an Applied Biosystems 7000 Real time system. Data was analyzed using the delta delta CT method. The following primers were used in the study:

~~~
Rpr2-F:TGGGTTGGCTCATGCTTATT
Rpr2-R:ATCCGAAGACCGGAAGAAAG
5kb_RA-F:CCGTCTACGGCCTTTGTTTA
5kb_RA-R:AGTGGAAGAACCAACCTGACA
5kb_lei-F:TTTTCGGAATGGGTTTTCAG
5kb_lei-R:ACACACACGAACCGAATGAA
GRIM2-F:TTATGCCAACAACCAACCAA
GRIM2-R:CCCCCTTTCTAGTTCCGAAG
AbdbChip2-F:TCTACTCCACCGGTTTGCTC
AbdbChip2-R:ACAGGCGGTCCTTATTGATG
intergenicSKB-F:TCAAGCCGAACCCTCTAAAAT
intergenicSKB-R:AACGCCAACAAACAGAAAATG
rpr_pro_F:AGAAGGCCAAAATGAGCAGC
rpr_pro_R:GCGCACACACTTTTCTTTCG
Act5C_f ATGTGTGTGTGAGAGAGCGA
Act5C_b AAACCGACTGAAAGTGGCTG
~~~

### Nuclear RNA preparation

After isolation of nuclei, as described above, proteins were digested and crosslinking reversed in 20mM Tris/1mM CaCl2/0.5%SDS with 1U/ul RNAse inhibitor and 500ug/ml proteinase K (Roche) at 55o for 3 hours. Approximately 600,000 nuclei were used for RNA purification with RNAzol. A mix of oligodT and random hexamer primers were used for reverse transcription, and quantitative PCR was done on an Applied Biosystems 7000 Real time system. DNA contamination was assesses with a no reverse transcriptase control, and data was analyzed using the delta delta CT method. The following primers were used for qPCR:

~~~
dRP49-F: 5′ CTC ATG CAG AAC CGC GTT TA 3′
dRP49-R: 5′ ACA AAT GTG TAT TCC GAC CA 3′
SA-F: GGACAAGATAATACCACCCGC
SA-R: CGCTTGATCAGTTTCGCCAT
~~~

### Generation of CRISPR deletion

Small deletions (120-130bp) of the Cut binding site upstream of the *rpr* promoter were generated with CRISPR/Cas9 (Fig. S4). Two guide RNAs were cloned in pCFD4 Vector (Addgene, 42749411) as previously described (Port et al., 2014). Stable gRNA transgenics were made by BestGene (CA, USA) and crossed with nos–cas9 (BL#54591). The progeny were screened for deletions by PCR, and breakpoints were confirmed by sequencing. Two lines CRISPR_ILB_2.9 and 431CRISPR_ILB_3.1 were used for phenotypic analysis.

### Fly stocks and genotypes

All the Flies were raised at 25 °C. Wild-type fly lines used in this study are yw^67c23^ and wor-Gal4/+. The following lines were obtained from the stocks centers at Bloomington, IN, USA (BL) and Vienna, Austria (VDRC) and by personal communications: repo-Gal4 (BL), UAS-nls-dsRed (BL), abdA-RNAi (BL) and cut-RNAi(BL, VDRC), SA RNAi (BL), NippedB RNAi (BL) UAS-Cut::UAS-mcd8-GFP (BL, Norbert Perrimon), nos–cas9 (BL), *cut*^c145^ (BL), cut-RNAi; cut-RNAi (provided by Y. N. Jan), UAS-abdA.HA (provided by Y Graba). The enh1-GFP transgenic line was previously generated in our lab (Arya et al 2015).

### Immunostaining and Fluorescent in situ hybridization (FISH)

Staining of whole embryos and larval CNS were done as described previously (Arya et al., 2015). The following primary antibodies were used in various combinations: rat anti-Dpn (1:150, Abcam, Cambridge, MA, USA), goat anti-AbdA (1: 500, dH17, Santa Cruz, CA, USA), rabbit or mouse anti-GFP antibody (1: 1000, rabbit, Invitrogen, Grand Island, NY, USA or mouse jl8, Clontech, Mountain View, CA, USA), mouse anti-Cut (1:700, DSHB, Iowa City, IA, USA), rabbit anti-H3K27me3 (1:500, Active Motif, 39157), rabbit anti-H3K9me3 (1:500, Abcam, ab8898), rabbit anti-H3K27ac (1:500, Active Motif, 39136), rabbit anti-H3K4me3 (1:500, Active Motif, 39159), and rabbit anti-Stromalin (1:700, a gift from Dale Dorsett). Secondary antibodies (Molecular Probes, Eugene, OR, USA) were used at 1:200 dilution.

FISH was performed as previously described (Arya et al., 2015). Digoxigenin (DIG)-labeled probes for *grim*, *rpr*, and GFP were used. When expression levels are compared, in situs were processed in parallel, and imaged with matched confocal settings and image processing. Embryos were imaged with any Nikon A1SiR confocal (Melville, NY, USA). Image Processing was done using Nikon Elements, ImageJ or Adobe Photoshop.

The intensity of histone marks was quantified with ImageJ software: average fluorescence intensity within a hemisegment of the nervous system was calculated and normalized to the intensity of a corresponding area within the epidermis. This calculation was performed across 11 confocal slices beginning with the ventral-most slice of the nervous system, as determined by Deadpan staining. Data in Fig. S6 are presented as the ratio of the average intensity of signal within the nervous system to the average intensity of signal within the epidermis at a given confocal slice.

SA levels in NBs and neurons were quantified in ImageJ. SA intensity was measured in 10 NBs or neurons in A3 to A6 segments of 3 embryos. To account for embryo-to-embryo variability in staining, SA levels in NBs or neurons was normalized to the average SA intensity outside of nervous system in the same stack.

## Acknowledgments

We would like to thank Norbert Perrimon, Y-N Jan and the Bloomington stock center for fly stocks, Dale Dorsett and the Developmental Studies Hybridoma Bank for antibodies, Sarah Bowman, Katia Georgopoulos and Georgopoulos lab members for advice on ChIP studies and data analysis, the White lab and Katia Georgopoulos for comments on the manuscript. This work was funded by R01GM110477 (KW & LZ), R03NS092063 (KW) and an MGH Fund for Medical Discovery award (RA) and R01GM106174 (LZ).

---

**Supplementary Figure 1.**
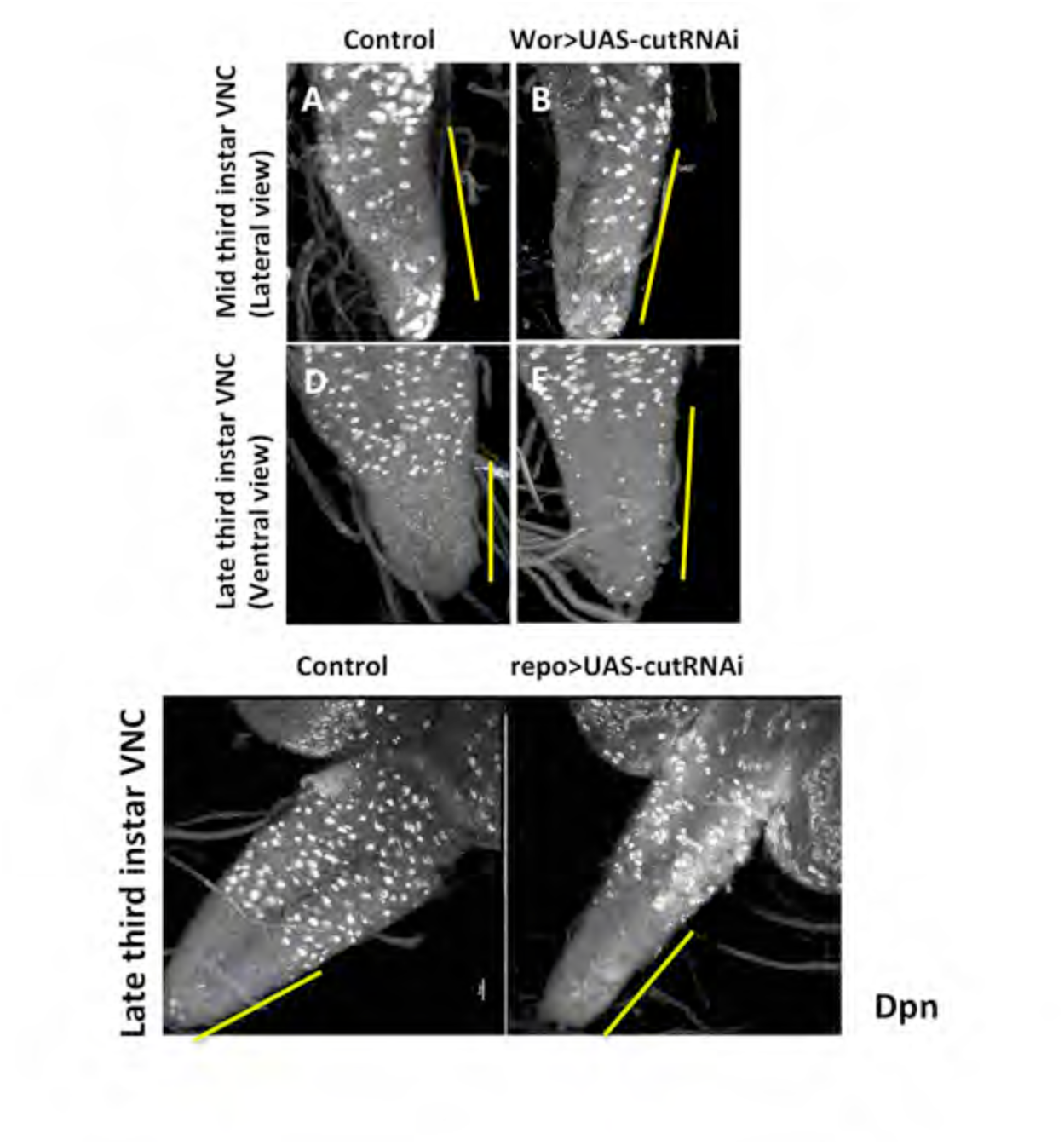
Knockdown of *cut* in the CNS with wor>cutRNAi results in ectopic NB survival in larvae. In mid-third instar larvae, ectopic NBs are clearly visible in the abdominal ganglia. In late third instar larvae, many of these extopic NBs have died, but at least one ectopic NB lineage remains. Knockdown of *cut* in glia with repo>cutRNAi does not result in ectopic NB survival.

**Supplementary Figure 2.**
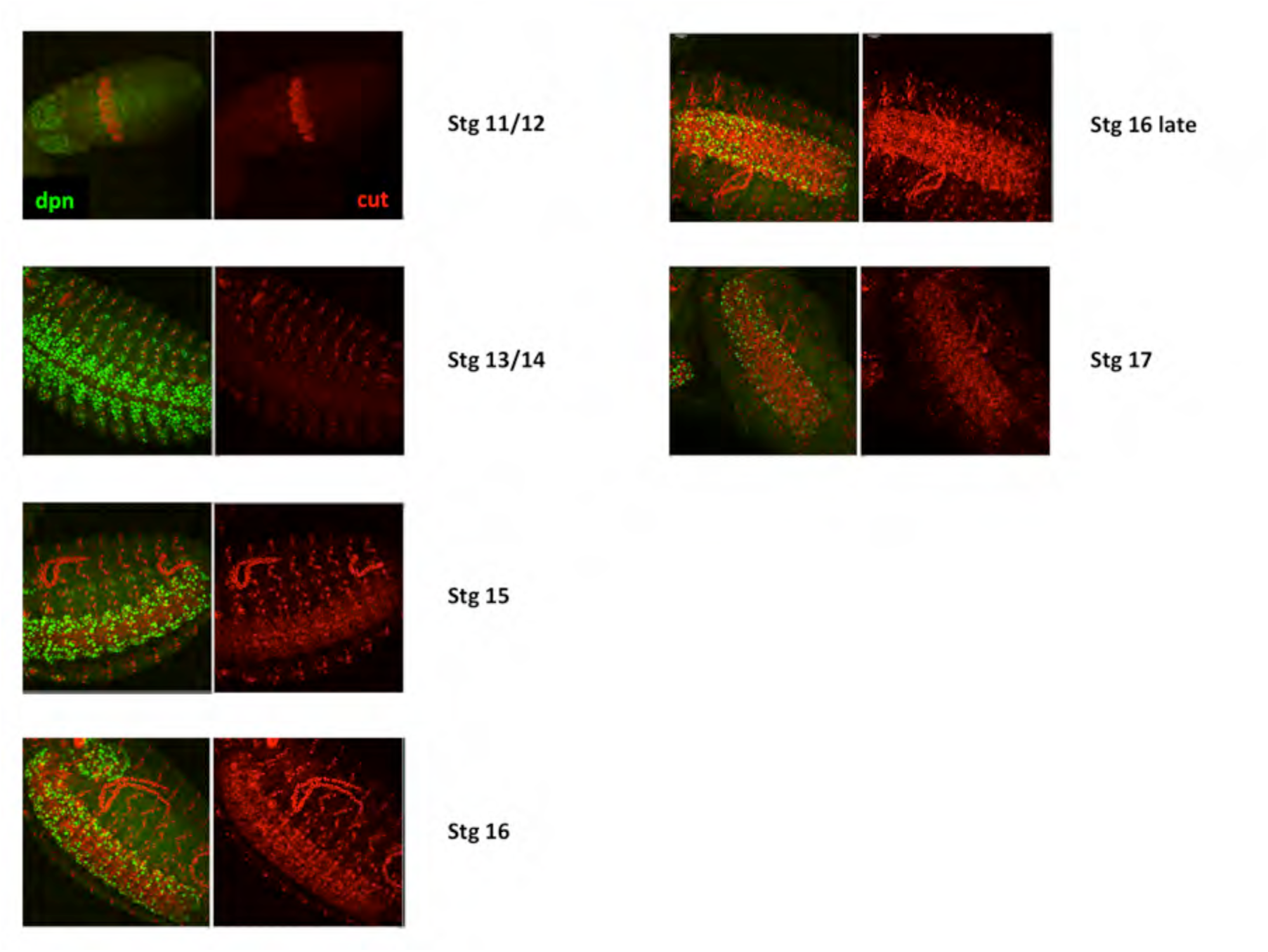
Cut expression in the CNS corresponds to the period of NB death. Cut levels increase in the CNS starting at stage 13/14, and are highest by late stage 16. All embryos were imaged at the same settings for Cut staining intensity. Dpn staining marks the NBs.

**Supplementary Figure 3.**
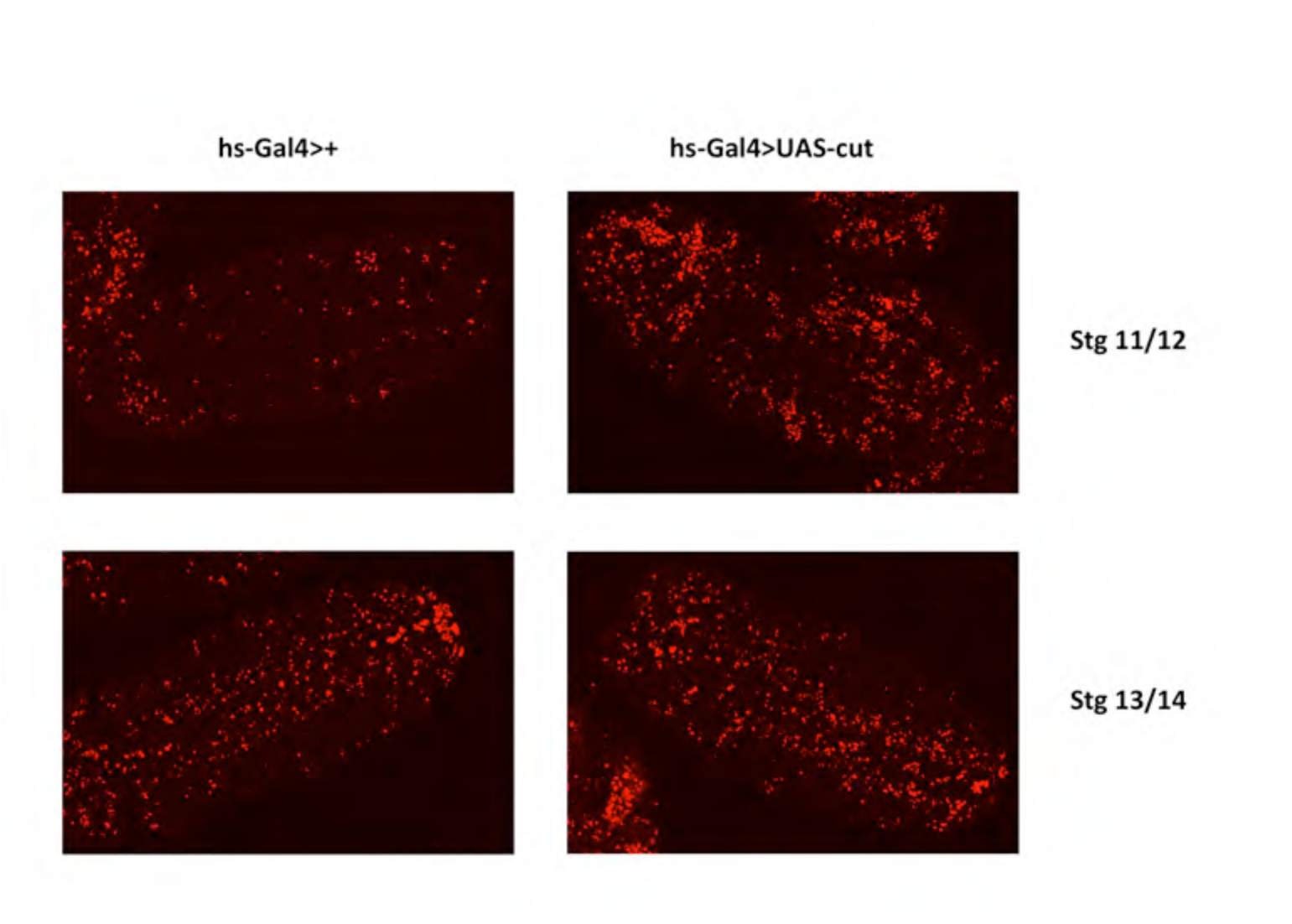
Expression of *cut* under control of the heat shock promoter results in a large increase in cell death, both within the nervous system and in other tissues. TUNEL staining marks apoptotic cells. An overnight collection of embryos from the cross hs-gal4 X UAS-cut mcd8GFP/Bal was heat shocked for 1.5 hours at 37o, allowed to recover for 4 hours and then stained for TUNEL and GFP, GFP staining is not shown.

**Supplementary Figure 4.**
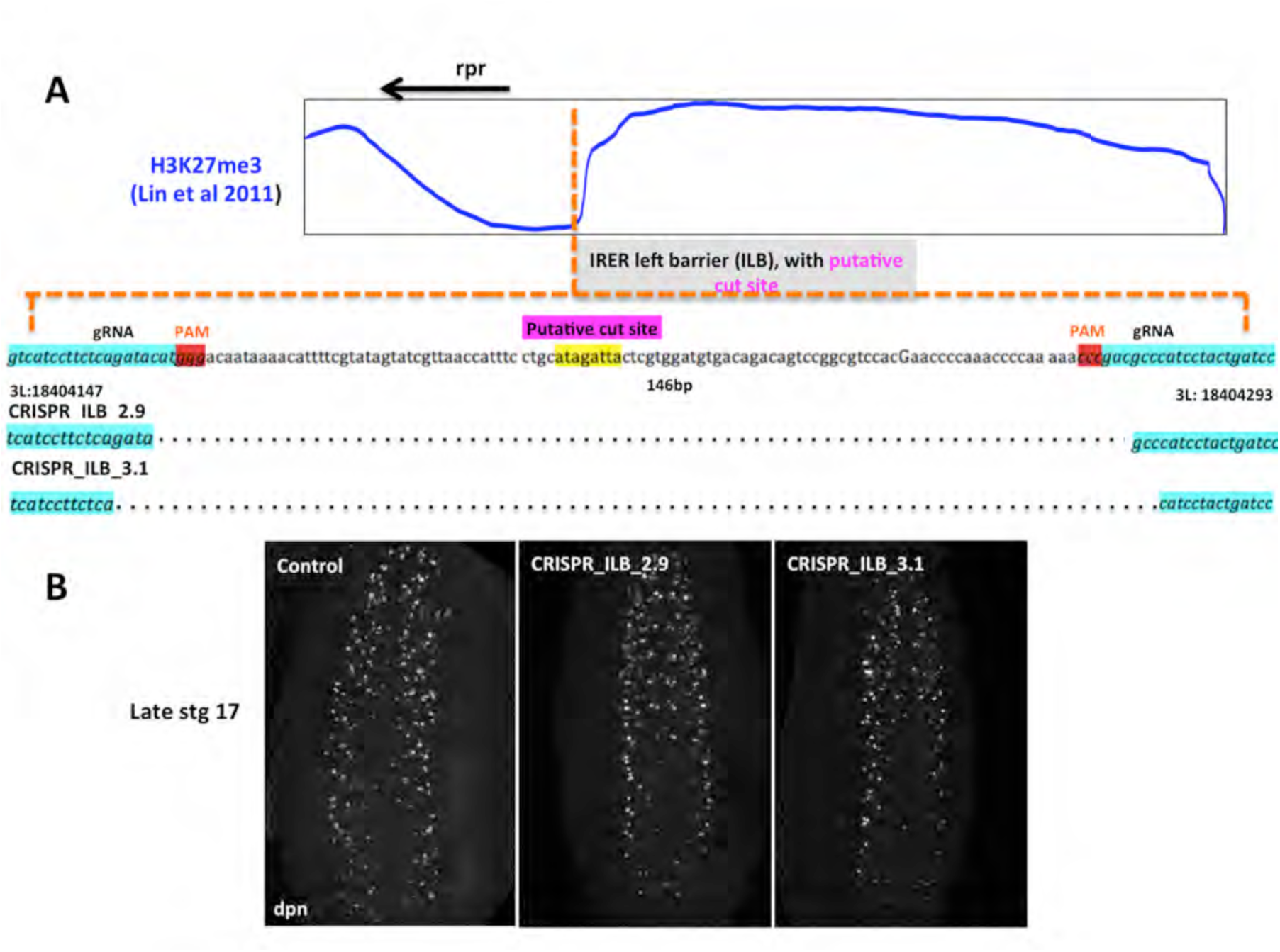
Crispr strategy for deletion of the putative Cut binding site at the ILB. **A)** The reported Cut binding site lies within the IRER left boundary, 5’ to the *rpr* transcribed region (Lin et al., 2011). Guide RNAs for CRISPR/Cas9 were selected to flank the binding site. Transgenic gRNA flies were crossed to nos-Cas9, and screened for deletions by PCR. Breakpoints were confirmed by sequencing. B) Late embryos homozygous for two deletions of the Cut binding site do not result in ectopic NB survival.

**Figure S5.**
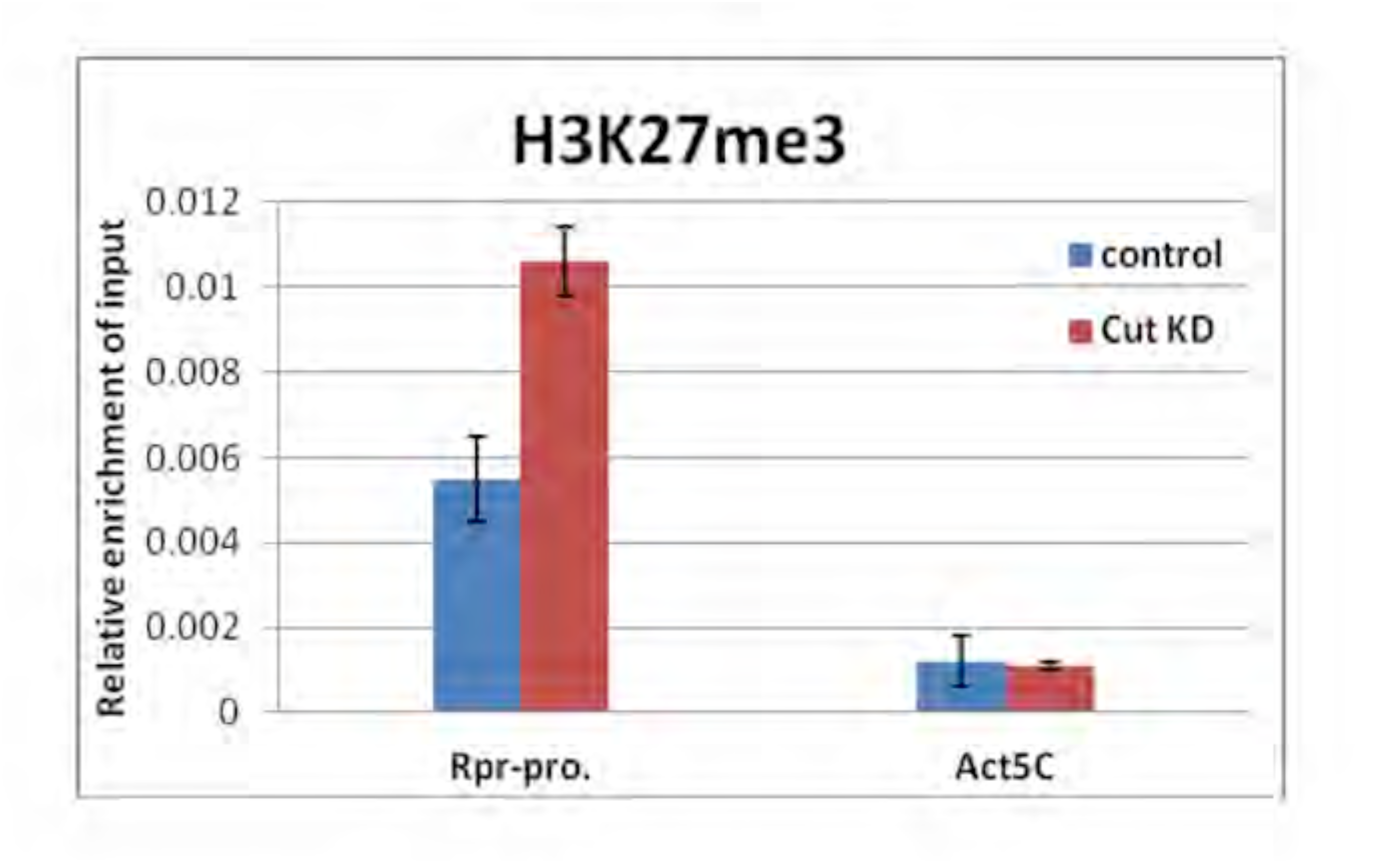
H3K27me3 enrichment at the rpr promoter as demonstrated by qPCR. ChIP with an independent chromatin preparation from sorted nuclei demonstrates increased enrichment of H3K27me3 at the *rpr* promoter following *cut* knockdown with wor>cutRNAi.

**Figure S6.**
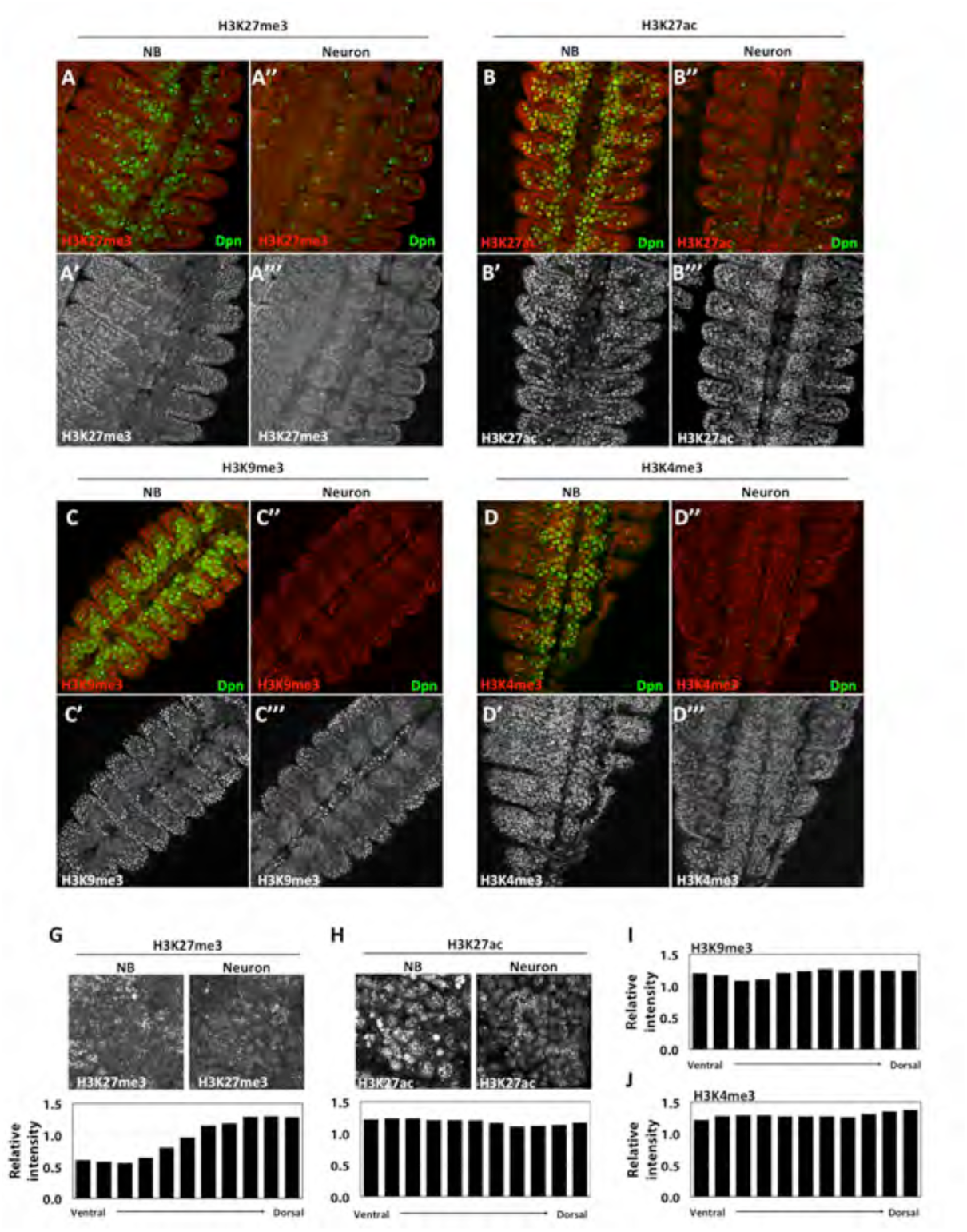
H3K27me3, but not other histone modifications, is differentially expressed between neuroblast and neuron layers of the ventral nerve cord (VNC) of stage 13 embryos. H3K27me3 (**A-A’’’**) is absent from Dpn-positive neuroblasts (**A** and **A’**) but is present in dorsal layers of the nerve cord that contain neurons (**A’’** and **A’’’**). H3K27ac (**B-B’’’**), H3K9me3 (**C-C’’’**) and H3K4me3 (**D-D’’’**) are present in both neuroblast and neuronal layers of the VNC. Relative H3K27me3 levels increase along the ventral-dorsal axis in the VNC: H3K27me3 staining is shown within one hemisegment of the VNC at the level of neuroblasts (**E, top left**) or neurons (**E, top right**), and quantified as relative intensity within each single confocal slice, normalized to the epidermis (**E, bottom**). In contrast, there is no change in relative H3K27ac levels along the ventral-dorsal axis, as shown in one hemisegment at the level of neuroblasts (**F, top left**) or neurons (**F, top right**), and quantified as in E (**F, bottom**). There is also no change in relative levels of H3K4me3 in the CNS, as shown in **H. G)** H3K9me3 is generally lower in the VNC overall, and shows slight reduction in relative levels in NBs at this stage.

**Figure S7.**
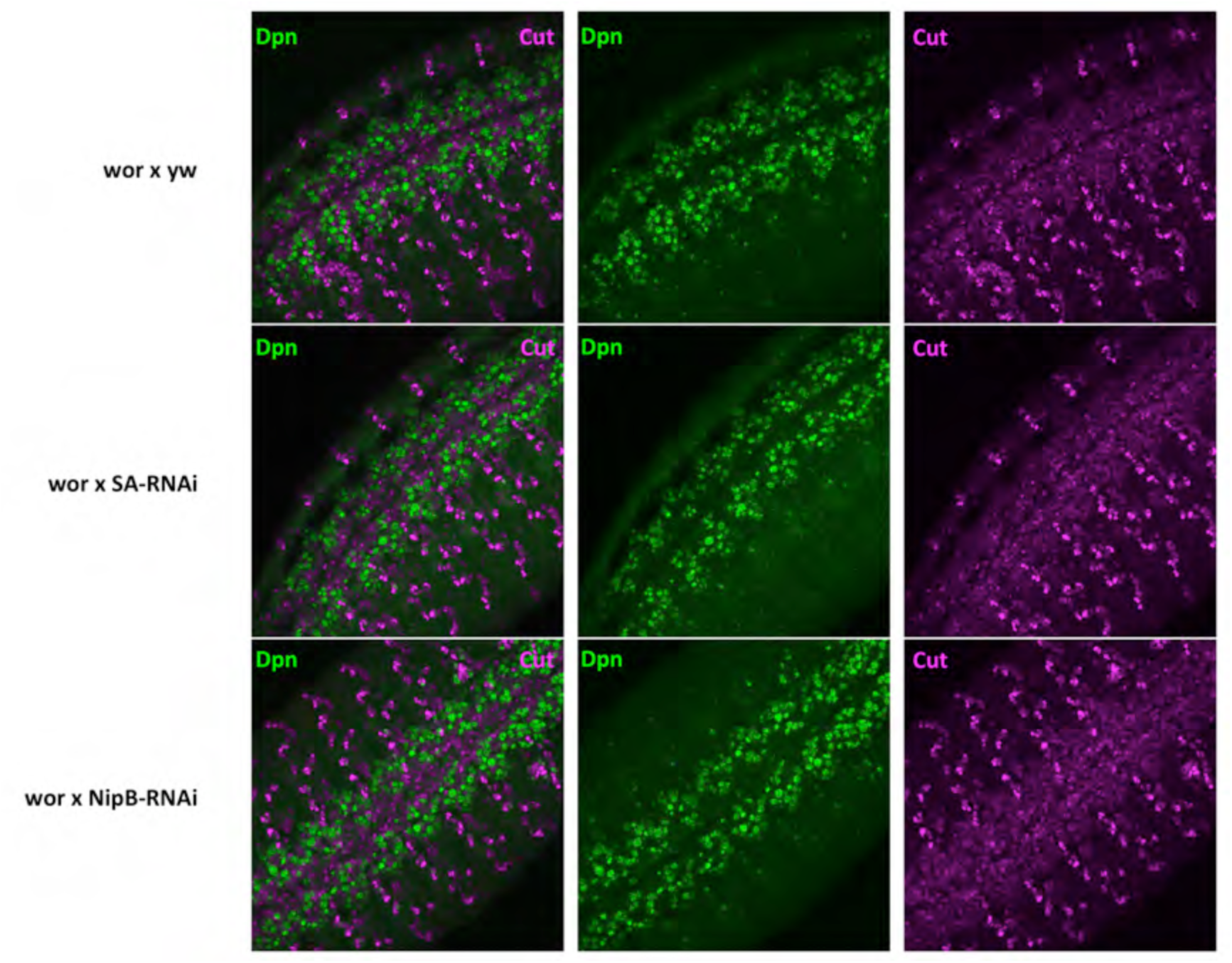
SA and Nipped-B do not regulate Cut expression in the CNS. Embryos expressing RNAi against *SA* or Nipped-B under control of wor-gal4 were collected overnight, fixed and stained for Dpn and Cut. Stage 14 embryos from each genotype were imaged at the same settings. Maximum projections are shown.

